# The latitudinal gradient in hand-wing-index: global patterns and predictors of wing morphology in birds

**DOI:** 10.1101/816603

**Authors:** Catherine Sheard, Montague H. C. Neate-Clegg, Nico Alioravainen, Samuel E. I. Jones, Claire Vincent, Hannah E. A. MacGregor, Tom P. Bregman, Santiago Claramunt, Joseph A. Tobias

## Abstract

An organism’s ability to disperse influences many fundamental processes in ecology. However, standardised estimates of dispersal ability are rarely available, and thus the patterns and drivers of broad-scale variation in dispersal ability remain unclear. Here we present a global dataset of avian hand-wing index (HWI), an estimate of wingtip pointedness widely adopted as a proxy for flight efficiency and dispersal in birds. We show that HWI is correlated with geography and ecology across 10,391 (>99 %) bird species, increasing at higher latitudes and in migratory and/or non-territorial species. After controlling for these effects, the strongest predictor of HWI is temperature variability (i.e. seasonality), with secondary effects of diet and habitat type. Our analyses (1) reveal a prominent latitudinal gradient in HWI shaped by ecological and environmental factors, and (2) provide a global index of avian dispersal ability for wider use in community ecology, macroecology, and macroevolution.

**Statement of authorship:** The study was conceived by CS and JAT. Data collection was led by JAT, SC, and CS, with contributions from CS, MNC, NA, SEIJ, CV, HEAM, TPB, and SC. CS performed the analyses. CS and JAT wrote the manuscript and all authors revised the text.

## Introduction

Dispersal plays a key role in ecological processes at a range of spatial and temporal scales (Greenwood 1980; Bowler & Benton 2005). The likelihood or rate of dispersal by organisms is thought to influence macroevolutionary patterns of speciation and extinction (Mayr 1963; Kisel & Barraclough 2010; Smith *et al.* 2014) as well as macroecological patterns of geographical range (Gaston 2003; Lester *et al.* 2007; Fritz *et al.* 2012) and range overlap (Pigot & Tobias 2015; Pigot *et al.* 2018), and is thus a major factor driving community assembly (MacArthur & Wilson 1967). Within populations, dispersal is a critical component of meta-population and meta-community dynamics (Hanski 1998; Venail *et al.* 2008; Jacob *et al.* 2019), thereby regulating fundamental ecological processes including nutrient transfer, pollination, and seed dispersal (Viana Duarte *et al.* 2016). Variation in dispersal ability is also a critical factor in predicting biological invasions (Capinha *et al.* 2015), as well as species sensitivity to climate (Travis *et al.* 2013) and land-use change (Lees & Peres 2009; Bregman *et al.* 2014). Despite this broad relevance, however, we still know remarkably little about phylogenetic and geographical variation in dispersal ability and its underlying drivers (Dieckmann *et al.* 1999; Pigot & Tobias 2015).

Dispersal distances vary widely in vertebrate animals from small-scale movements in sedentary species to global journeys spanning both hemispheres in migratory species. On the one hand, this variation is thought to be largely driven by environmental factors such as climate. In particular, temporal variability in climate or resources is expected to favour increased mobility (Greenwood 1980; Dieckmann *et al.* 1999; Bowler & Benton 2005), which in turn is expected to generate a latitudinal gradient in dispersal ability because of increased seasonality towards the poles (e.g. Salisbury *et al*. 2012, Bregman *et al*. 2014). On the other hand, dispersal is expected to be linked to a range of species attributes only weakly correlated with latitude, including body size (Paradis *et al.* 1998), diet (Peach *et al.* 2001), and resource defence strategy (Forero *et al.* 1999). Estimating the relative roles of these environmental or ecological factors has proved challenging, however, because dispersal is difficult to quantify directly in most natural systems, particularly in a standardised way across large numbers of species (Dieckmann *et al.* 1999; Dawideit *et al.* 2009; Alzate *et al.* 2019).

Standard methods for quantifying dispersal, such as mark-recapture, GPS tracking, and estimates of gene flow, are time-consuming, expensive, and difficult to scale. Even in vertebrate animals, comparative studies of dispersal have been limited to very small sample sizes, typically in well-studied organisms and regions. For example, natal and breeding dispersal data were made available for 75 British bird species based on nearly 100 years of intensive mark-recapture data (Paradis *et al.* 1998), while a recent survey of mammalian movement was based on GPS data from only 57 species (Tucker *et al*. 2018). Until such measurements become easier to implement at a wide scale, the most promising approach for comparative analyses relies on standardised biometric indices of dispersal ability. Perhaps the most familiar of these indices, the hand-wing index (HWI), is a morphological metric linked to wing aspect ratio (Kipp 1959; Lockwood *et al.* 1998) and widely used as a single-parameter proxy of avian flight efficiency and dispersal ability (Weeks & Claramunt 2014; Stoddard *et al*. 2017; Pigot *et al*. 2018; Bitton & Graham 2015; Burney & Brumfield 2009; Kennedy *et al*. 2016; Chua *et al*. 2017; Claramunt *et al*. 2012). HWI provides an estimate of wing pointedness (Fig. 1a) and has become a mainstay of macroecological analyses partly because – unlike wing-aspect ratio, which must be measured on open wings - it can be calculated from measurements obtained from dried museum specimens (Fig. 1b). Current species sampling for HWI, however, remains taxonomically incomplete and biased towards temperate regions.

**Figure 1.**
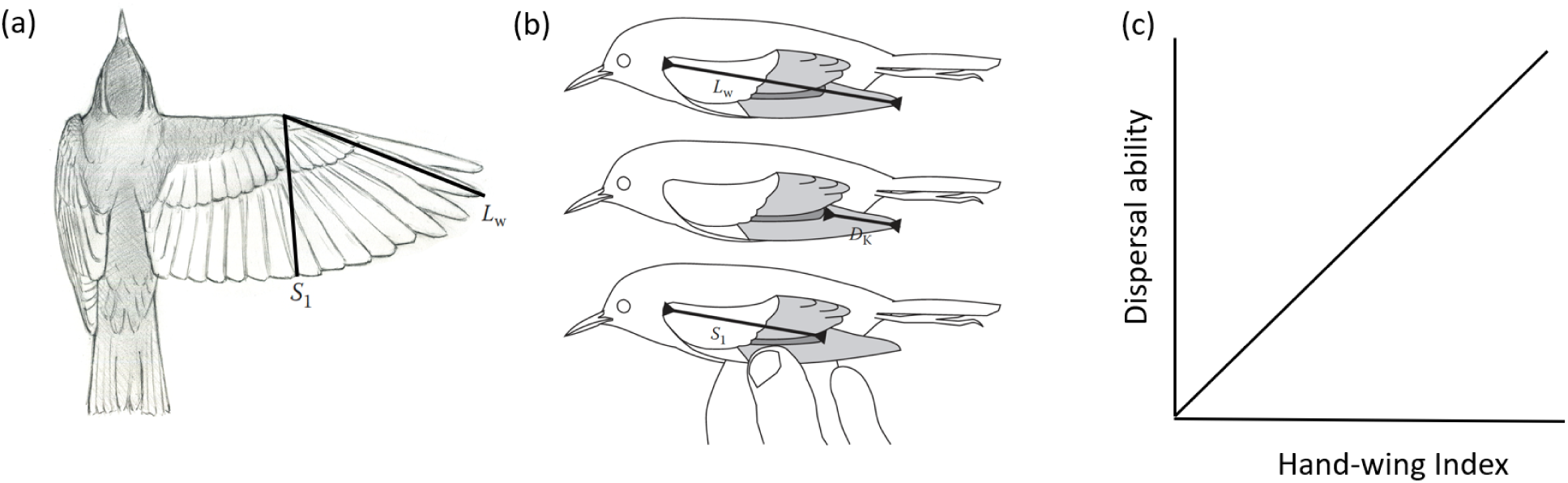
Calculating the hand-wing index (HWI), a proxy for wing aspect ratio, flight efficiency, and dispersal ability in birds. a, Open wing of passerine bird (Redwing *Turdus iliacus*, illustrated by Richard Johnson) showing wing length (*L*_w_) and secondary length (*S*_1_) used to generate HWI. b, Diagrams (reproduced from Claramunt & Wright 2017) showing these linear measurements taken on standard museum study skin. The first secondary is highlighted in dark grey. Kipp’s distance, D_K_, is defined as the difference between *L*_w_ and *S*_1_. c, HWI is an estimate of wing-aspect ratio, and in theory is positively associated with dispersal ability and flight efficiency (including flight strength, gap-crossing ability, and to a lesser extent natal and lifetime dispersal distance).

To provide a global synthesis of variation in avian HWI, we directly measured the wing morphology of 41,981 museum specimens and live birds representing 10,391 (>99%) bird species. For each species, we calculated average HWI from a combination of linear wing measurements, then mapped phylogenetic and spatial variation in HWI to investigate the global inter-specific drivers of avian wing morphology. Birds – the largest tetrapod radiation - provide an ideal test case for a broad-scale analysis of dispersal morphology because they are globally distributed and reasonably well studied, with a full species-level phylogeny (Jetz *et al.* 2012) and complementary datasets on geographical distribution, along with a range of ecological, social and other life history variables (Wilman *et al.* 2014; Tobias *et al.* 2016; Tobias & Pigot 2019).

We first use spatial mapping to visualise the geographic distribution of HWI, and then apply Bayesian phylogenetic mixed models to explore the mechanisms underlying this pattern. We include two biogeographic, four climatic, and five ecological variables to assess which of these factors best explain interspecific variation in HWI, both across all birds and separately within major groups (passerines versus non-passerines). We also test the hypothesis that wing morphology drives variation in geographical range size and use separate models to further explore the link between HWI and migration. Given that a growing body of evidence suggests that HWI predicts flight efficiency and dispersal ability in birds (Fig. 1c) (Marchetti *et al.* 1995; Lockwood *et al.* 1998; Dawideit *et al.* 2009; Baldwin *et al.* 2010; Claramunt & Wright 2017), our results provide insight into the factors shaping the evolution of dispersal across larger spatial and temporal scales and the consequences of dispersal for widely observed biogeographic patterns.

## Materials and Methods

### Data collection: HWI

The hand-wing index, HWI, is defined as the ratio of the Kipp’s distance (the distance between the tip of the first secondary feather and the tip of the longest primary feather) to the total wing chord (see Fig. 1a,b). HWI can therefore be viewed as Kipp’s distance corrected for size. High HWI indicates pointed (high aspect ratio) wings suitable for efficient long-distance flight, whereas low HWI indicates rounded (low aspect ratio) wings associated with weaker or short-distance flight (Claramunt *et al.* 2012; Weeks & Claramunt 2014; Stoddard *et al.* 2017). Kipp’s distance and HWI have been found to relate to inter-specific variation in natal dispersal distance and migration in a sample of British birds (Dawideit *et al.* 2009), as well as to flight ability and migratory behaviour more generally in passerines (Lockwood *et al.* 1998; Baldwin *et al.* 2010; Claramunt *et al.* 2012; Claramunt & Wright 2017). Flightless birds have lowest HWI (Stoddard *et al*. 2017).

We measured Kipp’s distance and the length of the unflattened wing chord to the nearest 0.5 mm in 41,981 museum specimens and live birds representing 10,391 extant and recently-extinct species. To allow global-scale phylogenetic analyses, we assigned species limits according to the Global Bird Tree (Jetz *et al*. 2012), adopting recent taxonomic revisions or newly described species where possible. We selected at least two males and two females in good condition for measurement when available, giving a mean of 4.04 individuals measured per species. We restricted our sample to adults, excluding all specimens labelled or identified in-hand as juveniles. Measurements from museum specimens were taken from the nominate subspecies whenever possible.

The majority (68%) of measurements were taken by 8 observers (the authors), but a further 84 observers contributed to the dataset by providing measurements from specimens accessed at a total of 73 collections and field sites (see Supplementary Information). To maximise consistency, all contributors were supplied with a detailed protocol for taking wing measurements. To assess the effect of observer biases, we collected 220 replicate wing measurements by different measurers for 146 species (see Supplementary Information). Measurer identity explained approximately 0.5% of the variation in Kipp’s distance and less than 0.01% of the variation in wing chord (Fig. S1). These findings support previous analyses concluding that independent biometric trait measurements by different observers are very highly correlated and thus unlikely to influence multi-species analyses at macroecological scales (McEntee *et al.* 2018).

### Data collection: ecological and social predictors of dispersal

We classified all species into dietary guilds using proportional diet scores from Wilman *et al.* (2014), updated according to the procedures described therein with information from primary and secondary literature (e.g. Birds of North America Online and HBW Alive). We merged all four vertivore categories (including carnivores and piscivores) into a single grouping. Any species with > 50% of its diet belonging to a single category, or with exactly 50% of its diet in one category and < 50% in all other categories, was classified as belonging to that majority category as a guild; all other species were classified as omnivores.

Territorial behaviour was classified into a binary score based on data obtained from Tobias *et al.* (2016): species were treated as territorial if they had year-round (‘strong’) territoriality, and non-territorial if they either lacked any form of territorial behaviour or if they were previously classified as having diffuse or seasonal (‘weak’) territoriality, including defence of nest-sites and mating display sites. Habitat use was also classified into a binary score according to data in Tobias et al. (2016): closed habitats include forests and semi-open habitats such as parkland, shrubland and marsh vegetation; open habitats include grasslands, deserts, coasts and oceans. Migration data was obtained from BirdLife International (2018), with ‘full migrants’ scored here as migratory and all others (partial migrants, altitudinal migrants, non-migrants, and nomads) scored as non-migratory. Body mass data was updated from Dunning (2007) based on primary and secondary literature (Tobias & Pigot 2019).

We computed geographical range size by intersecting global range polygons (Birdlife International & NatureServe 2011) with a 1°×1° grid and counting the number of grid cells overlapped by each polygon. These ranges were then intersected with data from WorldClim v. 1 to obtain average annual temperature, temperature variability, annual precipitation, and precipitation variability for all species (Hijmans *et al.* 2005). We used a land GIS layer to quantify the proportion of the geographical range of each species intersecting with islands with landmass below 2,000 sq. km (see Pigot *et al*. 2018), which we term ‘association with islands’. We extracted median range latitude from range polygons and entered this into models as an absolute (unidirectional) value. For data display purposes, range maps were intersected with a 1°x1° grid (approximately 110 km x 110 km), and the traits of the species living within each grid cell were plotted using the R package sf (Pebesma 2018).

### Bayesian phylogenetic mixed models

Bayesian phylogenetic generalised linear mixed models were run using the R package MCMCglmm (Hadfield 2010). We tested the effect of multiple variables – body mass, migration, diet, territoriality, latitude, association with islands, and climatic factors – on mean species HWI, with phylogeny as a random effect. All quantitative variables were scaled to have a mean of 0 and a variance of 1; range size and HWI were log-transformed; and the association with islands was arcsine-transformed. Priors were initially set using inverse-Wishart priors for the phylogenetic and residual variance (V = 1, ν = 0.002) and diffuse normal priors for the fixed effects (mean 0, variance 10^10^). After conducting a dummy run of 11,000 iterations on an arbitrary tree with a burn-in of 1,000 and a thin of 50 to determine a start point for the R- and G-structures, each of 100 tree topologies was run sequentially for 15,000 iterations with a burn-in of 5,000 and a thin of 1,000, for a total posterior sample of 1,000 solutions (10 per tree). All chains were visually inspected to ensure proper mixing, and autocorrelation was checked using the command ‘autocorr’ with 0.1 used as a target threshold. These models were run separately for all birds, passerines, and non-passerines.

We also ran an additional set of models testing the predictive effects of HWI on geographical range size and migratory strategy. Range size is likely a consequence, rather than an explicit driver, of wing morphology, so we tested whether HWI influences the extent of the global distribution of bird species using a model including migration as a potential covariate. The extent to which wing morphology promotes migration is less obvious, particularly as HWI seems to be highly labile. High HWI is often rapidly lost in sedentary or insular taxa (Kennedy *et al.* 2016; Hosner *et al.* 2017), suggesting that it can also readily evolve in lineages that switch to a migratory strategy. Nonetheless, high HWI may predict migration if wings adapted for functions such as dispersal or foraging facilitate the switch to long-distance dispersal strategies. We therefore tested whether wing morphology explains long-distance migration, with other potential drivers held constant. These models included an additional interaction effect of absolute latitude and hemisphere, allowing separate coefficients to be modelled for the HWI-migration relationship in the northern or southern hemispheres. As migratory strategy is here considered a binary variable, we used a logistic regression (MCMCglmm family ‘categorical’) with priors for the fixed effects set using the command ‘gelman.prior’, an improper prior for the phylogenetic variance (V = 10^−10^, ν = −1), and the residual variance fixed at 1. All phylogenetic analyses were run on a sample of 100 trees obtained from the Hackett backbone of the Global Bird Tree (Jetz *et al.* 2012).

## Results

### Patterns of variation in HWI

Across all birds, HWI ranges from 0.016 in *Rhea pennata* to 74.8 in *Phaethornis ruber* (and is arbitrarily set as 0.1 in the five wingless kiwi species, Apterygiformes). At the scale of clades, HWI is lowest for the ratites (e.g. Struthioniformes, mean 0.019) and highest for the tropicbirds (Phaethontiformes, mean 69.2) (Fig. 2). Viewing total phylogenetic variation in this trait reveals that, on average, HWI is lower across passerines than across non-passerine orders (Fig. 2). Within both major clades, notable peaks in HWI coincide with highly dispersive clades such as parrots (Psittacidae), pigeons (Columbidae), shorebirds (Charadriiformes), seabirds, and waterfowl, as well as groups specialised on foraging in flight, such as swallows (Hirundinidae) or swifts and hummingbirds (Apodiformes).

**Figure 2.**
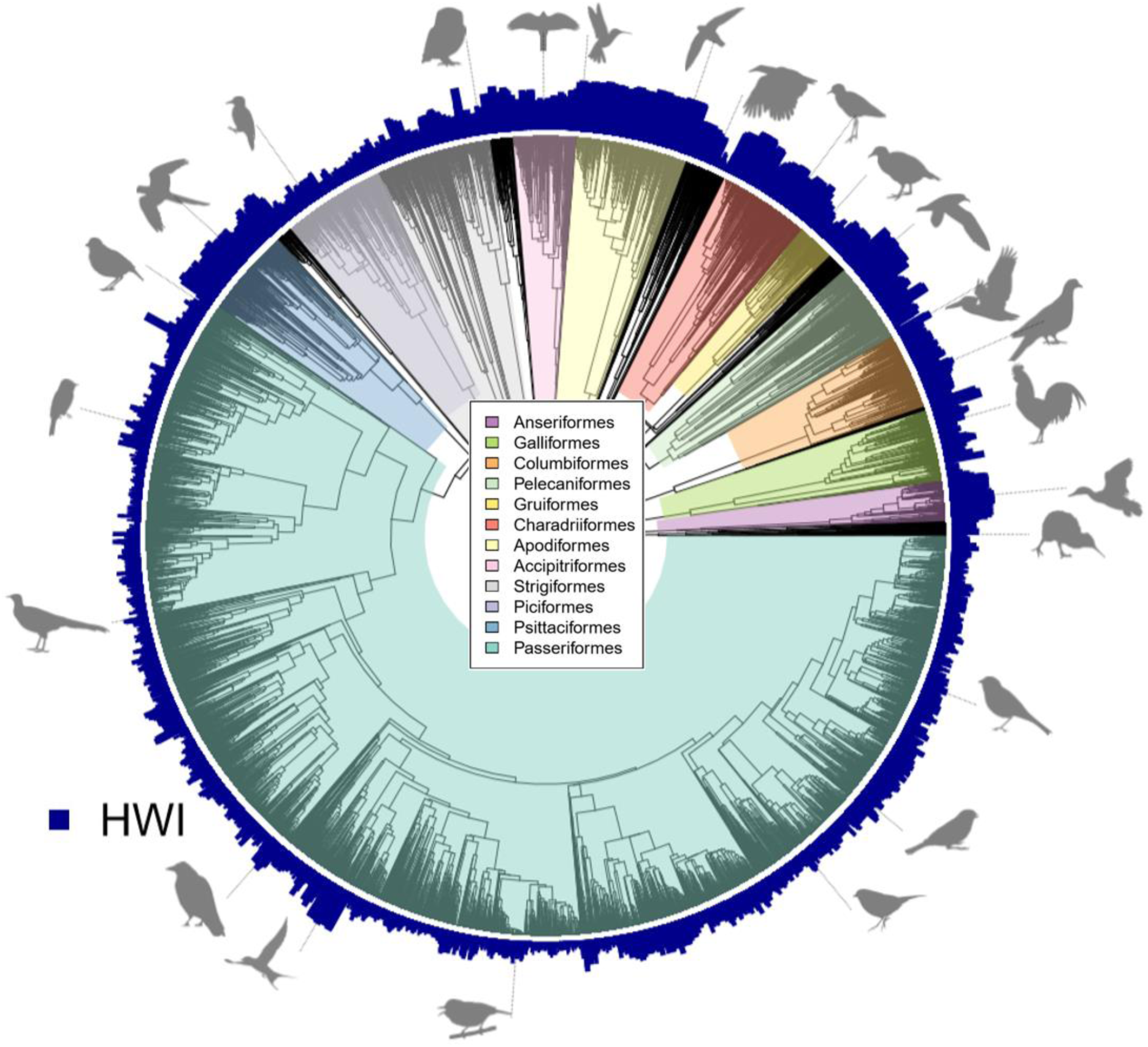
The phylogenetic distribution of hand-wing index (HWI) values across birds (n = 9,945 species). Variation in HWI is plotted at branch tips of a single phylogenetic tree extracted from www.birdtree.org (Jetz *et al*. 2012) using the Hackett backbone. For ease of interpretation, the 12 most species-rich clades are highlighted. Silhouettes are from *phylopic.org*, with full attribution information in the supplementary materials.

Focusing on geographical trait variation (Fig. 3), we find that the highest average HWIs are found in scattered regions worldwide, notably in the high Arctic, and in drylands such as the Saharan and Arabian deserts. In addition, a pronounced latitudinal gradient in HWI is clearly visible, with the lowest values consistently found in tropical regions. These spatial patterns are largely recapitulated in both non-passerines and passerines although the latitudinal gradient is shallower in passerines with relatively high HWI much more broadly distributed across the temperate zone, particularly in the northern hemisphere. This apparent relationship between HWI and certain biomes suggests that avian flight capacity is broadly related to climatic conditions, in particular environmental variability or unpredictability.

**Figure 3.**
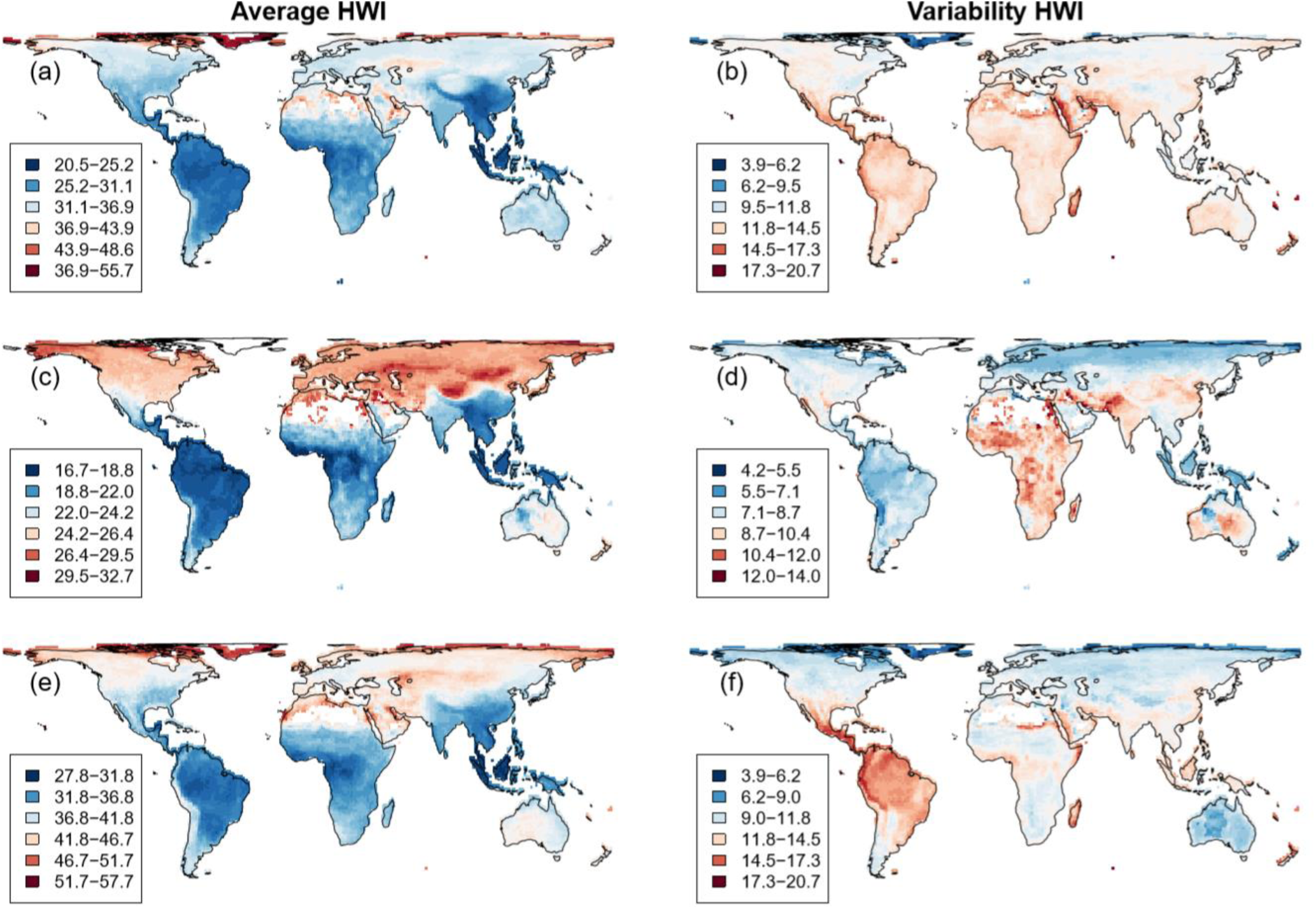
Global variation in hand-wing index (HWI) for all birds (a, b; n = 8,504), passerines (c, d; n = 5,153), and non-passerines (e, f; n = 3,351). HWI is calculated for grid-cell assemblages as both average (a, c, e) and variability (b, d, f). Each grid cell represents a 1⁰×1⁰ square (approximately 110 km × 110 km).

Focusing on trait variability within assemblages (Fig. 3), we find that the highest variability in HWI is in the Saharan and Arabian deserts, the Andes mountains, Madagascar, and the Pacific islands (e.g. New Zealand, New Caledonia, Fiji, Hawaii, and the Galapagos). In other words, these are hotspots of variability supporting a wide spectrum of dispersal traits from low to high HWI. Different patterns emerge within major clades: variability in passerine HWI peaks in Africa, Australia, and south-central Asia from Iran to Pakistan, whereas in non-passerines variability peaks in New World low latitudes. Thus, the co-occurrence of different HWI values is driven not only by environment, but by the separate evolutionary and biogeographic histories of different clades.

### Ecological and environmental drivers of HWI

Across all birds, HWI is strongly positively correlated with breeding range temperature variability and in particular migration (Fig. 4). These two factors are themselves correlated because migration tends to arise in species breeding in highly seasonal environments (see Tables S4-6). Conversely, HWI is strongly negatively correlated with year-round territory defence, presumably because this behaviour is tightly bound to a relatively sedentary lifestyle, and is also strongly linked to tropical biomes, particularly tropical forests (Tobias *et al.* 2016). We find further significant, though weaker, correlations with diet, habitat type, and association with islands, as well as latitude and climate. Specifically, high HWI is associated with nectarivory, small body masses, open habitats, islands, high precipitation, high temperatures, and low latitudes. Conversely, species that eat invertebrates or that breed in locations with high precipitation variability are more likely to have low HWI. Although body mass is traditionally used as an index for dispersal in vertebrates, with larger size assumed to indicate greater dispersal ability (e.g. Sutherland et al. 2000; Whitmee & Orme 2013), we find a negative correlation between avian body mass and HWI. This result is not surprising given that some of the largest bird species (e.g. ratites including ostriches, rheas and cassowaries) are highly sedentary with low HWI.

**Figure 4.**
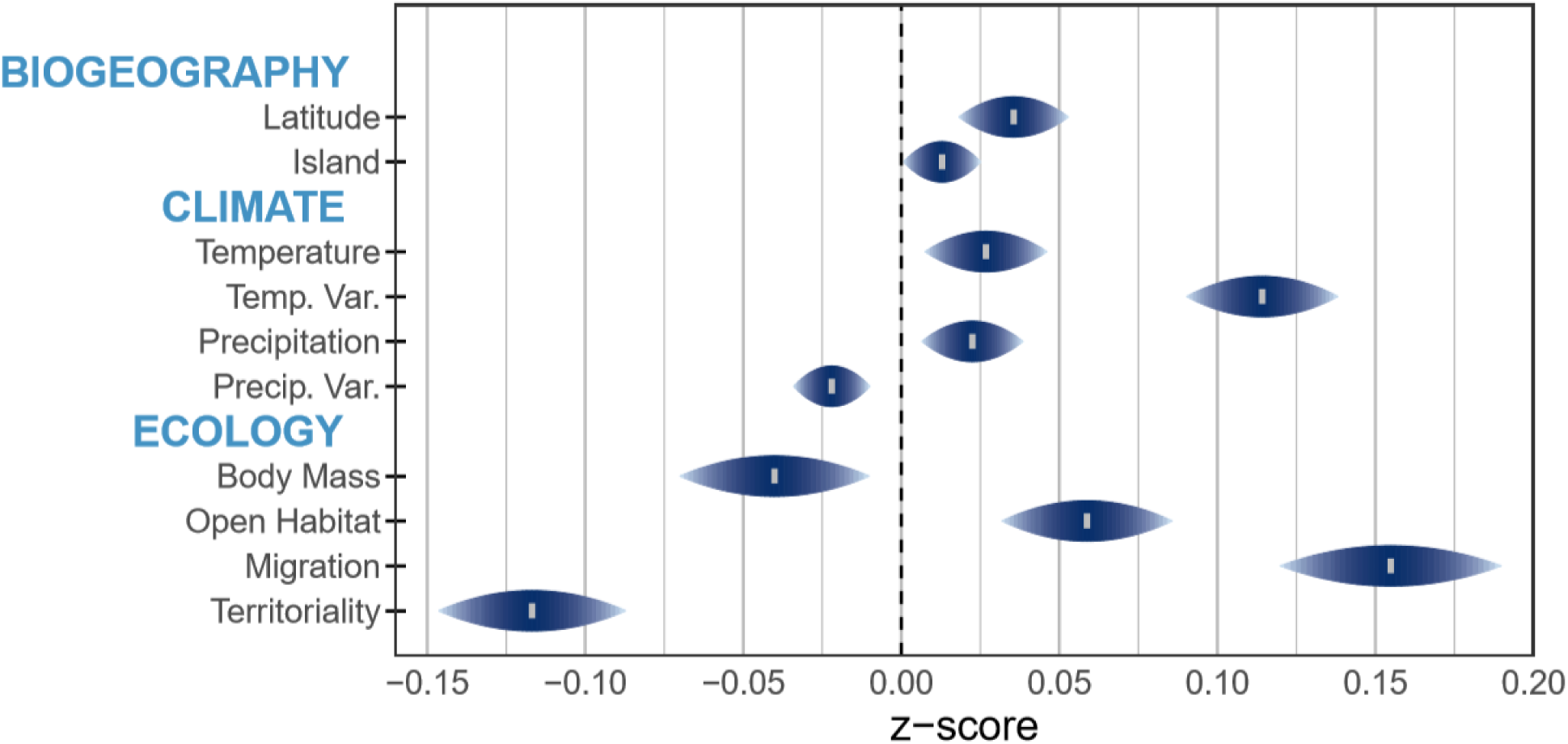
Predictors of hand-wing index (HWI) in birds (n = 9,277 species). Shown are z-scores and 95% credible intervals (CI) computed with Bayesian phylogenetic mixed models; the dashed line represents a coefficient of 0. High z-score indicates positive association with HWI; low z-score indicates negative association with HWI. Climatic variables are calculated for each 1⁰×1⁰ grid cell of geographical range and averaged. Temperature = annual mean temperature. Temp. Var. = variation in monthly temperature values over a year (standard deviation). Precipitation = annual precipitation. Precip. Var. = variation in monthly precipitation values over a year (coefficient of variance). Open Habitat = grasslands, deserts, coasts, oceans. Dietary categories are omitted for ease of interpretation.

**Figure 5.**
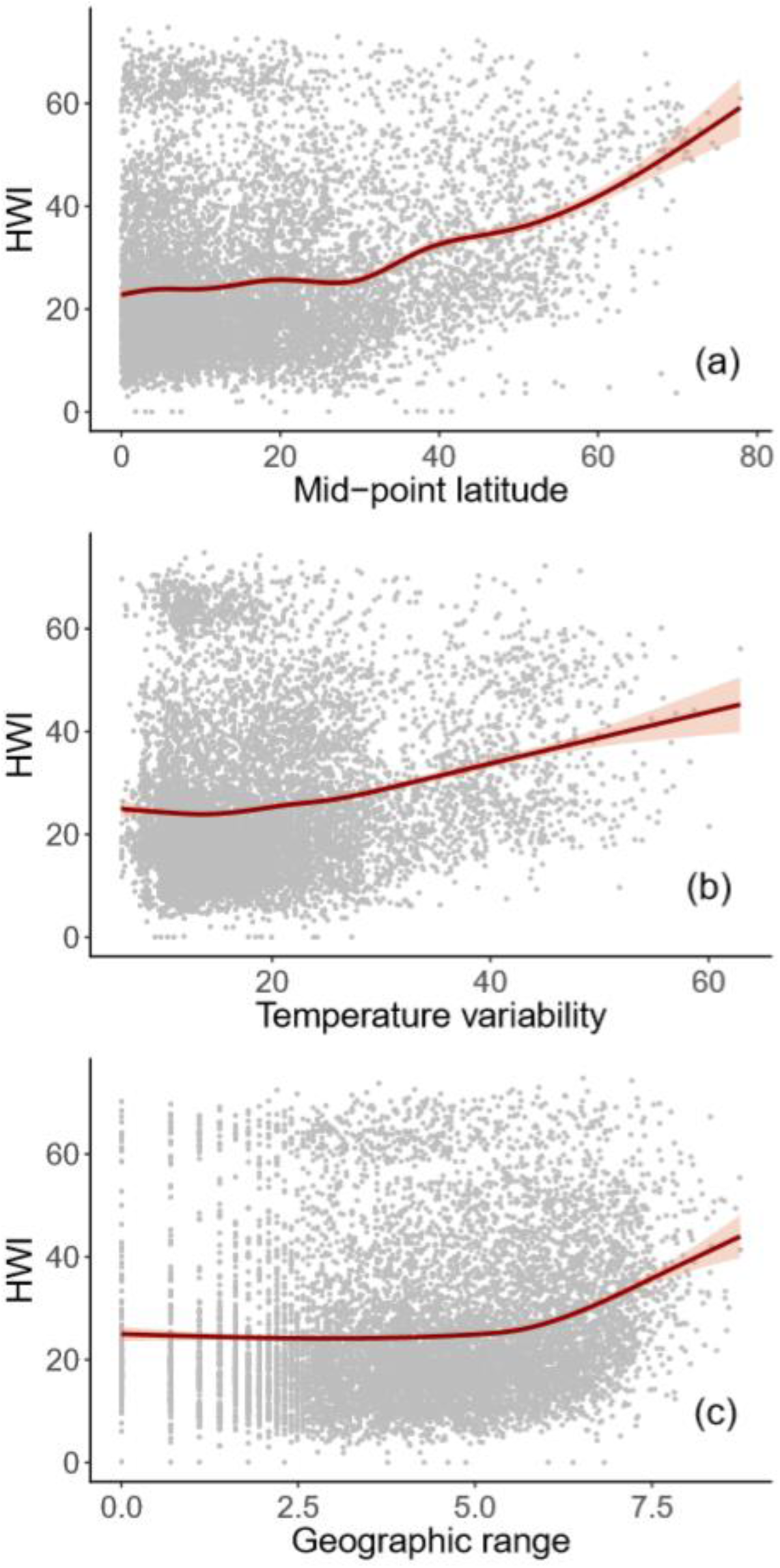
Relationship between hand-wing index (HWI) and environmental variables for birds (n = 9,356 species). Panels show how HWI varies with (a) latitude, (b) temperature variability, and (c) geographical range size. All environmental variables are generated from polygons of breeding ranges; climatic variables are calculated for each grid cell of the range and averaged; overall range sizes (km^2^) are log-transformed. White line shows model fit and red shading shows confidence intervals.

**Figure 6.**
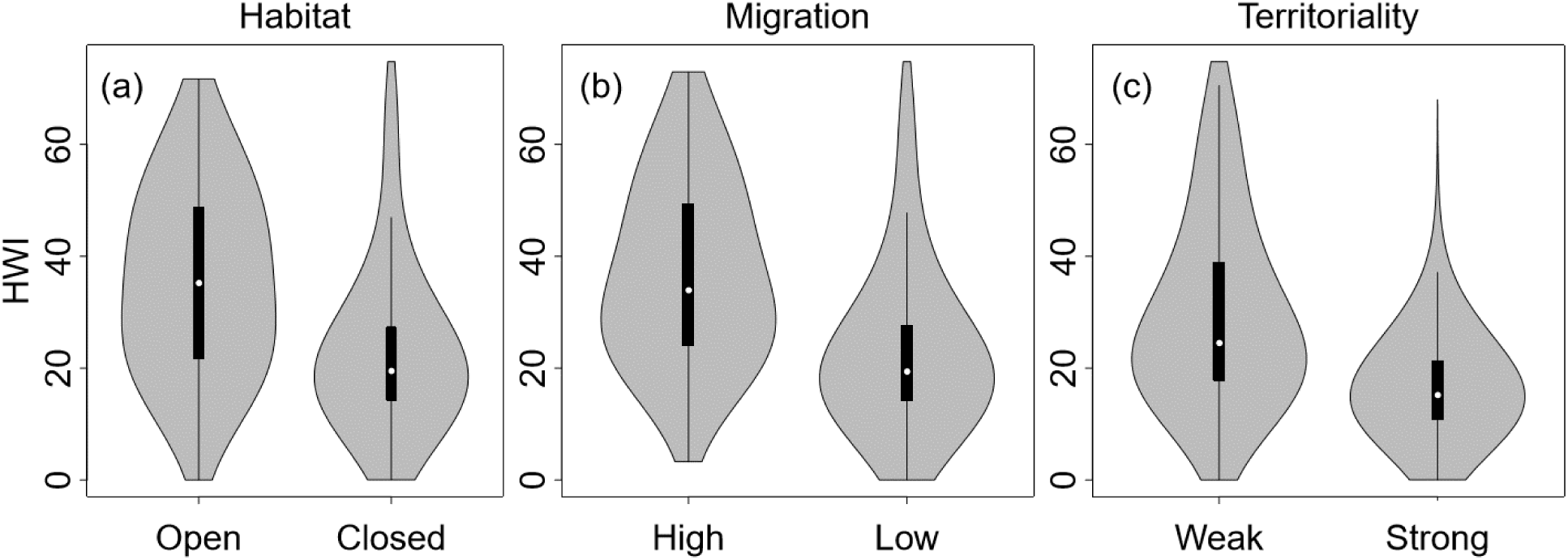
Relationship between hand-wing index (HWI) and ecological variables. A global sample of bird species (n = 9,854) were classified according to (a) primary habitat (Open = grassland, deserts, shrubland, parkland, thorn forest, seashores, cities; Closed = forests), (b) migration (High = >50% geographical range entirely vacated during non-breeding season; Low = sedentary, elevational migrants, partially migratory, or <50% geographical range migratory), and (c) territoriality (Weak = defending territories only seasonally or never holding territories except very small lek or nest-site territories; Strong = defending territories year-round). Black interior boxplots show median (white dot), first and third quartile (ends of black box), and data minimum and maximum excluding outliers (whiskers). The shaded kernel probability densities illustrate the distribution of the data.

Across passerines, the correlates of HWI are similar to those for all birds, although temperature and body mass are no longer significant predictors (see Fig. S2). In non-passerines, HWI is correlated with neither diet nor breeding range precipitation seasonality nor association with islands (see Fig. S3). The tendency within non-passerines for smaller species to have larger HWIs is perhaps strongly driven by swifts and hummingbirds (Apodiformes) being adapted for sustained flight whereas many larger non-passerines are flightless.

### HWI as a predictor of biogeography and migration

When we examined the biogeographic consequences of wing morphology, we found that multiple variables are correlated with range size in birds, namely temperature variability, precipitation, diet, latitude, migration, association with islands, and whether the species is found in the northern or southern hemisphere; there was also a weaker but still significant effect of habitat type and an interaction between latitude and hemisphere (Table S1). Some of these relationships reflect the well-established tendency for range size to increase with latitude (Rapoport’s rule) coupled with the much larger available land-area in the northern hemisphere. Accounting for all these factors, we found that species with higher HWIs have larger range sizes (z-score: 0.116, p < 0.001). The findings are qualitatively similar when the model is restricted to non-passerines or passerines (Tables S2-3).

HWI is the strongest predictor of migration across all birds (z-score: 2.268, p < 0.001), as well as in non-passerines and passerines, separately. The only other predictors strongly and positively associated with migration across all taxonomic categories are temperature variability and latitude (Table S4). In contrast with range size models, there are differences in secondary predictors of migration between non-passerines and passerines, including factors such as diet, climate and hemisphere (Table S5-6). It is difficult to infer causal mechanisms from these patterns because wing morphology could be a consequence of rather than a constraint on long-distance dispersal⎯indeed, specifically because HWI co-evolves with migration (Kennedy *et al.* 2016; Hosner *et al.* 2017). Either way, our analysis highlights the strong link between wing morphology and migration reported previously in birds (Lockwood *et al.* 1998; Dawideit *et al.* 2009).

## Discussion

Our global analysis of avian wing morphology reveals strong gradients in HWI shaped by environmental and ecological variables. We find that increased HWI is linked to both migration and territoriality, as well as high variability in breeding-range temperature, with habitat type and diet as secondary effects. Given that HWI is a standard proxy for flight strength and efficiency in birds (Dawideit *et al.* 2009; Baldwin *et al.* 2010; Claramunt *et al.* 2012), our results suggest that average dispersal ability declines in bird species that live in tropical regions. Likely contributing to this finding, tropical species often have sedentary lifestyles and/or year-round territoriality, both of which are promoted by low variability in breeding range temperature. These broad-scale variations in dispersal may contribute to a number of pervasive macroecological and macroevolutionary patterns, including latitudinal gradients in diversification rate (Cardillo *et al.* 2005), species richness (Mittelbach *et al.* 2007), and range size (Tomašových *et al.* 2016), as well as the heightened sensitivity to environmental change reported for tropical taxa (Stratford & Robinson 2005; Tewksbury *et al.* 2008; Bregman *et al.* 2014; Freeman *et al.* 2018).

Previous studies mapping inter-specific variation in dispersal ability have shown few consistent geographical trends in dispersal indices. This may reflect poor data quality and coverage, with the relevant indices either limited to indirect factors such as body size, or applicable to only a relatively small subsets of species or regions (e.g. Gaston & Blackburn 2003; Sekar *et al*. 2012; Kennedy *et al*. 2016; Tucker *et al.* 2018; Chen *et al*. 2019; Donati *et al*. 2019). In comparison with other dispersal-traits, HWI is more amenable to global sampling and has a stronger mechanistic link to flight ability through its association with wing aspect ratio (Lockwood *et al.* 1998). Whether HWI is also strongly associated with average dispersal distance for each species requires further empirical testing, but high HWI is likely associated with increased gap-crossing ability even in cases of sedentary tropical species (e.g. parrots, pigeons, hummingbirds). Our dataset thus provides unique insight into the broad-scale distribution of dispersal abilities in birds, at least until direct dispersal estimates can be sampled with standardised methods across thousands more species.

### Environmental variability and the evolution of dispersal

Most of the key predictors of variation in HWI, including year-round territoriality and migration, vary with latitude. In turn, these underlying gradients may be driven by the striking contrasts in climatic variability – or seasonality – towards the poles. It is clear that latitude itself is not the key factor driving dispersal because temperature variability is one of the strongest predictors of HWI, with an effect over double that of latitude. Temperature variability appears to be fundamental because neither temperature nor precipitation alone has a significant association with wing morphology. Conversely, in passerines, variability in precipitation has a negative relationship with HWI, perhaps because sedentary species abound in tropical regions with comparatively stable temperature regimes but pronounced wet seasons.

A link between temporal climatic variability and increased dispersal has been demonstrated in a variety of taxa (Greenwood & Harvey 1982; Bowler & Benton 2005), in line with theoretical predictions (Dieckmann *et al.* 1999; Jocque *et al.* 2010). Our findings in relation to temperature variability support the prevailing view that species inhabiting stable environments are more likely to be sedentary, thus lacking dispersal-related adaptations, whereas other species inhabiting uncertain or highly seasonal environments can thrive in these conditions by relocating in space, often over long distances.

### Ecological drivers

Several ecological variables may influence variation in dispersal traits among species, potentially mediating the link with climate. The most obvious factor is year-round territoriality, a resource-defence strategy associated with reduced HWI in our analyses. Since year-round territoriality has an effect similar in strength and direction to that of low migration, our models suggest that territorial behaviour influences wing morphology above and beyond its association with migration: in both migratory and non-migratory species, those that hold year-round territories have lower HWI than those that do not. These findings are consistent with ecological theory predicting that individuals or species defending resources will have lower rates of dispersal compared with non-territorial individuals or species (Greenwood 1980; Greenwood & Harvey 1982; Clarke *et al.* 1997; Bowler & Benton 2005).

Diet is another factor partially associated with latitude or environmental variability, although the underlying relationships with HWI are complex. For example, previous studies have suggested that insectivory is associated with larger dispersal distances than other trophic niches (Peach *et al.* 2001), reflecting the traditional focus on temperate-zone systems where insectivorous birds tend to be migratory in response to seasonal prey availability. In contrast, tropical insectivores are mostly sedentary, and often year-round territorial (Salisbury *et al*. 2012), which explains why low-latitude insectivores have lower HWI than most other dietary niches in tropical systems. Nectarivores, by contrast, have the highest average HWI whether viewed across all birds or only in passerines. Moreover, nectarivore HWI does not tend to decline towards the tropics, presumably because of behavioural aspects of the foraging niche (Tobias & Pigot 2019). Strong flight ability makes sense in this guild because many nectarivores either forage on the wing or move across large areas to exploit patchy and unpredictable resources (Warrick 1998). Specialist nectarivores are largely confined to the tropics, yet this reverse trend of high HWI in a tropical guild is obfuscated at global scales because nectarivores are outnumbered by other trophic niches (e.g. insectivores, granivores and omnivores) that have relatively low HWI in the tropics.

### Linking HWI with gradients in biodiversity and conservation

According to speciation theory, dispersal is a key factor determining the likelihood of speciation in a given geographical area because it influences rates of gene flow across barriers (Kisel & Barraclough 2010). On islands, speciation is most likely to occur when dispersal is high enough to promote island colonisation (Fritz *et al.* 2012; Kennedy *et al.* 2016) but not so high that gene flow is too frequent (Diamond *et al.* 1976; Weeks & Claramunt 2014). On continents, however, most evidence suggests that low dispersal promotes avian diversification (e.g. Burney & Brumfield 2009; Claramunt *et al*. 2012; Cadena *et al*. 2019). Even on larger islands, such as Borneo, low dispersal (inferred from low HWI) strongly predicts reduced gene flow and incipient speciation in birds (Chua *et al.* 2017). Thus, our finding of a latitudinal gradient in HWI is consistent with the view that dispersal constraints promote allopatric speciation in tropical birds, contributing to the latitudinal diversity gradient (Salisbury et al. 2012).

Dispersal has also been linked to geographical range size and range overlap. For example, it is often assumed that strong dispersal facilitates colonisation of new regions, thereby driving range expansion, although empirical evidence linking dispersal and range size is largely inconclusive (Gaston 2003; Lester *et al.* 2007; Alzate *et al.* 2019). Our global analyses show a positive, though weak, correlation between avian HWI and range size, matching the findings of previous studies focused in different taxonomic groups (Luo *et al.* 2019) or at smaller taxonomic scales (Kennedy *et al.* 2016). Some studies also find evidence that variation in avian HWI predicts species coexistence (Pigot & Tobias 2015), with the initial establishment of range overlaps occurring more readily among lineages with higher HWI (Pigot *et al.* 2018). If avian dispersal ability increases with latitude, as suggested by spatial patterns in HWI, then this could help to explain why the same trend from tropics to poles arises in both range size (Rapoport’s rule; Tomašových *et al*. 2016) and the incidence of early sympatry (Martin *et al.* 2010).

Finally, global variation in HWI may highlight species and communities at risk from climate or land-use change. For example, our findings support previous suggestions that gap- or barrier-crossing ability declines towards the equator (Moore *et al.* 2008; Bregman *et al.* 2014) and that many tropical birds are therefore more susceptible to habitat fragmentation (Stratford & Robinson 2005; Tobias *et al.* 2013; Bregman *et al.* 2014). HWI can also be used to study the role of dispersal ability in determining species responses to habitat fragmentation and configuration (Habel *et al.* 2019). In addition, given the importance of dispersal as a key factor influencing how individual organisms and populations respond to environmental change, species-level variation in HWI may serve as a standardised trait for modelling historical biogeographic patterns (e.g. Sukumaran & Knowles 2018) as well as the effects of future climate and development scenarios on biodiversity (Travis *et al.* 2013; MacLean & Beissinger 2017; Tucker *et al.* 2018). Ongoing changes in landscape and climate are leading to increased distances between habitat patches, seasonal ranges, and migration stopovers (Howard *et al.* 2018). Ultimately, these future landscapes are predicted to select for greater dispersal ability, suggesting that adaptation may even cause gradients in HWI to shift or steepen over time.

### Conclusions

We have shown that three inter-connected factors – migration, territoriality, and environmental variability – are most strongly correlated with variation in wing pointedness (HWI), a standard index of dispersal ability in birds (Claramunt *et al.* 2012; Stoddard *et al.* 2017; Pigot *et al.* 2018). These findings suggest that environmental and behavioural factors act in tandem to shape biogeographic gradients in dispersal and confirm a prominent tendency towards reduced dispersal ability in tropical bird species. Although this latitudinal gradient in dispersal has received little attention, it may play a fundamental role in generating more familiar macroecological patterns. Specifically, our analyses highlight the potential roles of dispersal in driving latitudinal gradients of speciation, range size, and sensitivity to environmental change, and also provide a trait-based template for exploring these roles at a range of scales from local communities to global ecosystems.

## Acknowledgements

For access to specimens, we thank the many curators and collections managers listed in the Supplementary Information, especially Hein van Grouw, Mark Adams, and Robert Prys-Jones of the Natural History Museum, Tring. For help with data collection and curation, we thank Bianca Darski, Daniel Swindlehurst, Sarah Rosenberg-Wohl, Phil Chapman, and numerous other contributors listed in the supplementary material. We are grateful to Gavin Thomas and Rich Grenyer for comments on a previous version of this manuscript. This study was funded by Natural Environment Research Council grants (NE/I028068/1, NE/K016385/1) (to JAT), the Oxford Clarendon Fund (to CS), the US-UK Fulbright Commission (to CS), and the Natural Sciences and Engineering Research Council of Canada (NSERC) Discovery Grant RGPIN-2018-06747 (to SC).

